# Theoretical and empirical quantification of the accuracy of polygenic scores in ancestry divergent populations

**DOI:** 10.1101/2020.01.14.905927

**Authors:** Ying Wang, Jing Guo, Guiyan Ni, Jian Yang, Peter M. Visscher, Loic Yengo

## Abstract

Polygenic scores (PGS) have been widely used to predict complex traits and risk of diseases using variants identified from genome-wide association studies (GWASs). To date, most GWASs have been conducted in populations of European ancestry, which limits the use of GWAS-derived PGS in non-European populations. Here, we develop a new theory to predict the relative accuracy (RA, relative to the accuracy in populations of the same ancestry as the discovery population) of PGS across ancestries. We used simulations and real data from the UK Biobank to evaluate our results. We found across various simulation scenarios that the RA of PGS based on trait-associated SNPs can be predicted accurately from modelling linkage disequilibrium (LD), minor allele frequencies (MAF), cross-population correlations of SNP effect sizes and heritability. Altogether, we find that LD and MAF differences between ancestries explain alone up to ~70% of the loss of RA using European-based PGS in African ancestry for traits like body mass index and height. Our results suggest that causal variants underlying common genetic variation identified in European ancestry GWASs are mostly shared across continents.

---

Polygenic scores (PGS, also known as PRS when applied to diseases) are now routinely utilised to predict complex traits and risk of diseases from findings of genome-wide association studies (GWASs). Over recent years, the predictive performances of PGS have steadily increased with GWASs sample sizes, as predicted by theory^1^. However, the over-representation of European ancestry in the majority of GWASs has been shown to yield an unbalanced improvement of PGS prediction accuracy, in particular in non-European ancestry populations^2,3^. For example, Duncan et al.^2^ report the average accuracy of PGS across multiple traits to be ~64% lower in individuals of African ancestry as compared to individuals of European ancestry. Similarly, Martin et al.^3^ report, across multiple traits, reductions of PGS accuracy relative to European ancestry of ~37%, ~50% and ~78% in individuals of South-Asian, East-Asian and African ancestries respectively.

Although increasingly emphasised in the recent GWAS literature, it is worth noting that the “loss of accuracy” problem is not utterly new. Indeed, a number of studies in the animal breeding literature have previously reported lower accuracy of genomic selection across genetically distant breeds^4,5^, consistent with the observation of limited transferability of GWAS findings across diverse human populations^6,7^. These studies also highlight major factors influencing that loss such as differences between populations in causal variants effect sizes, in alleles frequencies and in linkage disequilibrium (LD) between causal variants and SNPs assayed in GWAS.^6,8,9^ To illustrate the latter point, consider a SNP which LD *r*^2^ with a causal variant equals 0.8 in the discovery population and 0.6 in the target population. Such a SNP would therefore explain 25%=(1-0.6/0.8) less trait variation and thus be less predictive in the target population as compared to the discovery population, even when causal variants and their effect sizes are shared between ancestries. More generally, previous empirical and simulation studies have shown that accuracy of genetic predictors decays monotonically with increased genetic differentiation (*F*_ST_) and LD differences between ancestries^4,7,10^. Other factors such as population specific causal variants^11,12^ or gene×environment interaction^13,14^ have also been implicated as potential explanations of the loss of PGS prediction accuracy.

In addition, deterministic formulas have been derived to predict the accuracy of genomic prediction across breeds as a function of population parameters (e.g. heritability, genetic correlation) and also using selection index theory^15,16^. However, these deterministic formulas mostly apply to best linear unbiased predictors^17^, which are not classically used in human studies. Consequently, a theoretical understanding of the trans-ethnic predictive capacities of standard PGS is still missing. Although the key factors causing the loss of accuracy have been enumerated in previous studies, quantification of their relative contributions has not been done systematically. Note that quantifying the relative contributions of all these factors is critical for understanding aetiological differences between ancestries, which may have important clinical implications. Here, we develop a new theory to predict the relative accuracy of PGS in an ancestry divergent sample as a function of population genetics parameters. Our method only requires GWAS summary statistics and ancestry specific reference panels. We evaluate the performances of our theory through extensive simulations and apply it to GWAS of height and body mass index (BMI) in ~350,000 unrelated UK Biobank participants.

## RESULTS

### Expected accuracy of PGS in ancestry divergent populations

We consider a quantitative trait *y,* for which the genetic component is underlain by random additive effects of *M_c_* causal variants. Without loss of generality, we assume causal variants to be shared between ancestries but allow their effect sizes to vary from one ancestry to another. Therefore, ancestry-specific causal variants are a special case with non-zero effect sizes in only one ancestry. We then assume that a GWAS of *y* has been performed in a discovery sample of a given ancestry, hereafter denoted by Population 1 and that a PGS, defined as the sum of minor allele counts weighted by their estimated effect sizes from the discovery GWAS, is used to predict *y* in a target sample of another ancestry, hereafter denoted by Population 2.

We derived the expected accuracy of such PGS in Population 2 (denoted by 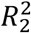) as function of the expected accuracy in a sample of same ancestry as Population 1 (denoted by 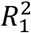), the minor allele frequency (MAF) *p_k,l_* at the *k*^th^ PGS-SNP, i.e. SNPs included in the PGS, in population *l,* the LD between the *j*^th^ causal SNP and the *k*^th^ PGS-SNP (denoted by *r_jk,l_*), the heritabilities 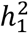 and 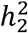 of *y* in Populations 1 and 2 respectively, and the correlation *r_b_* of causal SNP effect sizes between Population 1 and Population 2. Our main result is shown below in Equation (1) (details of our derivations are given in **Supplementary Note 1**):

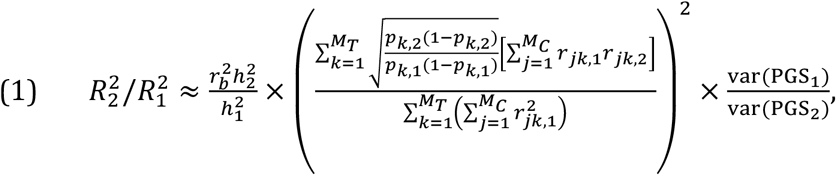

where *M_T_* denotes the number of GWS SNPs used to calculate PGSs.

Equation (1) shows that the relative accuracy (RA, relative to the accuracy in populations of same ancestry as population 1) of PGS, defined as 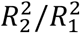, can be discomposed as the product of multiple terms: (i) the squared genetic correlation (the correlation of effect sizes of causal variants) between populations, (ii) the ratio of heritability between populations, (iii) the ratio of squared covariances between PGS and *y* in both populations; and (iv) the ratio of variance of PGS in both populations. This decomposition allows us to distinguish and thus separately quantify the fraction of the RA that is attributable to differences in effect size distributions between populations (term (i) and term (ii) in the decomposition, including 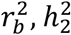 and 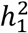) and the fraction attributable to alleles frequencies and LD differences between populations (term (iii) and term (iv) in the decomposition).

Many terms in Equation (1) can be quantified a priori using information from previous studies or from reference panels. However, the big unknown in Equation (1) remains the LD between unobserved causal variants and PGS-SNPs. Understanding how much PGS-SNPs tag causal variants is critical to quantify and therefore predict the RA of PGS. To reduce this uncertainty, we focus in this study on PGS based on independent genome-wide significant (GWS) SNPs, which are more informative of the location of causal variants than sub-significant SNPs. Importantly, previous studies^2,3,18,19^ have shown reduced predictive performance of PGS across ancestry when the PGS includes sub-significant SNPs, which provides an additional rationale for concentrating on GWS SNPs. Note also that in the near future, as GWAS sample sizes increase, the accuracy of GWS-based PGS will become similar to that of genome-wide PGS approaches.

Given that causal variants are largely unknown, we propose a heuristic method that considers as a “candidate causal variant”, any SNP in LD (*r*^2^>0.45) with a GWS SNP and located within 100 kb of the latter. This heuristics is justified by a previous study by Wu et al.^20^ which has quantified the fine-mapping precision of GWAS and has found over multiple computer simulations that causal variants lied within 100 kb of the GWS SNPs ~90% of the time and that LD *r*^2^ between causal and GWS SNPs was >0.45. Once candidate causal variants are identified for each independent GWS included in the PGS, we approximate Equation (1) by replacing 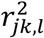 and *r*_*jk*,1_*r*_*jk*,2_ with the average of these quantities over all candidate causal variants, as shown below in Equation (2):

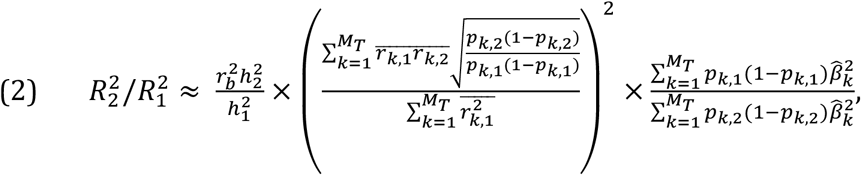

where 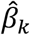 represents the effect size of the *k*^th^ PGS-SNP estimated in the discovery GWAS. We use the notation 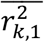 to denote the mean squared correlation of allele counts between the *k*^th^ PGS-SNP and all candidate causal SNPs within 100 kb. Similarly, we define 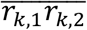 as the mean of *r*_*jk*,1_*r*_*jk*,2_’s between the *k*^th^ PGS-SNP and all candidate causal SNPs within 100 kb.

Moreover, we assume that the accuracy of the PGS in samples of same ancestry as the discovery GWAS is known (e.g. estimated in an independent sample) and propose to quantify other input parameters such as allele frequencies and LD correlations using data from an ethnically diverse reference panel like that of Phase 3 of the 1000 Genomes Project (1KGP)^21^. Finally, the quantification of *r_b_* and 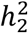 requires access to phenotypic and genotypic data from the target population, which may not be available simultaneously. Therefore, when such data are unavailable our method can only quantify the fraction of the RA that is explained by allele frequencies and LD differences between populations.

### Performance of the method on simulated data

We ran computer simulations to evaluate the performances of Equations (1) and (2) under various genetic architectures. We also assessed the performances of a naïve approach that assumes GWS SNPs to be the causal variants. In this case the expected RA explained by allele frequencies and LD differences between populations would approximately equal to 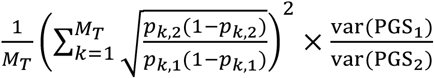. When PGS-SNPs are independent and given that SNP effect sizes from GWAS are typically small and of similar magnitude, this naïve approach can be further considered as a function of the ratio of heterozygosity at GWS SNPs between ancestries.

Our simulations utilise existing genotypes at ~1.1 million common HapMap 3 SNPs imputed in 351,983 unrelated UK Biobank (UKB) participants. These participants were categorised into four ethnically homogeneous groups corresponding to European ancestry (EUR; N_EUR_=333,263), East-Asian ancestry (EAS; N_EAS_=2,257), South-Asian ancestry (SAS, N_SAS_=9,448) and African ancestry (AFR; N_AFR_=7,015). The European ancestry group was further divided into a discovery set of N=313,284 participants in which GWS SNPs were identified (Online Methods), a validation set in which the accuracy of PGS within-European-ancestry was quantified and a reference group in which we predicted the accuracy of PGS. A thorough description of how these groups were defined is given in the Online Methods section. As our main focus is to predict the fraction of the RA that is attributable to alleles frequencies and LD differences between populations, we therefore assumed that effect sizes of causal variants are perfectly correlated across populations, i.e. *r_b_* = 1 and that heritability is constant across populations, i.e. 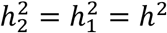. In total, we simulated six scenarios corresponding to three values of the number of causal variants *M_c_* = 1,000, 5,000 and 10,000; and two values of trait heritability *h*^2^ = 0.25 and 0.5 (Online Methods).

As expected, we observed across all scenarios that accuracies of PGS decreased monotonically with increased genetic distance to EUR (**Fig. S1**). The genetic distance was measured as *F*_ST_ described in **Supplementary Note 2.** More specifically, we found the largest RA in individuals of Asian ancestry (mean RA ~90.7% in SAS and mean RA ~76.6% in EAS), which has an average *F*_ST_ of ~0.06 with EUR. The smallest RA was observed in individuals of AFR (~46.4%), which has an average *F*_ST_ of ~0.14 with EUR (**Fig. S1 and Fig. S2**). These results imply that the predictive power of PGS remains limited even when causal variants and their effect sizes are shared between ancestries^19,22^. Over 100 simulation replicates (**Fig. 1**), we found that for all ancestries, the RA of PGS predicted using Equation (1) were not statistically different from the observed RA (*t*-test, *t*-statistic=0.68, *p*>0.05), which thus validates our theory. Moreover, we found that our heuristic approach based on candidate causal variants similarly yields unbiased predictions of the RA (*t*-test, *t*-statistic=1.34,*p*>0.05). However, the naïve approach assuming GWS SNPs to be the causal variants overestimated the RA in non-European ancestries (**Fig. 1**), with the overestimation ranging from 6.3% in SAS up to 84.5% in AFR. This result suggests that population differences in LD between causal variants and tag SNPs contribute a larger fraction to the decreased RA than allele frequencies differences at tag SNPs only. It is also worth noting that our predictions were nearly insensitive to either using whole-genome sequence (WGS) data from the 1KGP or imputed genotypes of UKB participants as reference panels (**Fig. S3**), which reflected by the highly correlated allele frequency (**Fig. S4**) and LD score (**Fig. S5**) between WGS and imputed data (**Supplementary Note 3**). Altogether, our simulation results show the validity of our theory and highlight its ability to accurately predict the RA attributable to allele frequencies and LD differences between ancestries.

**Figure 1.**
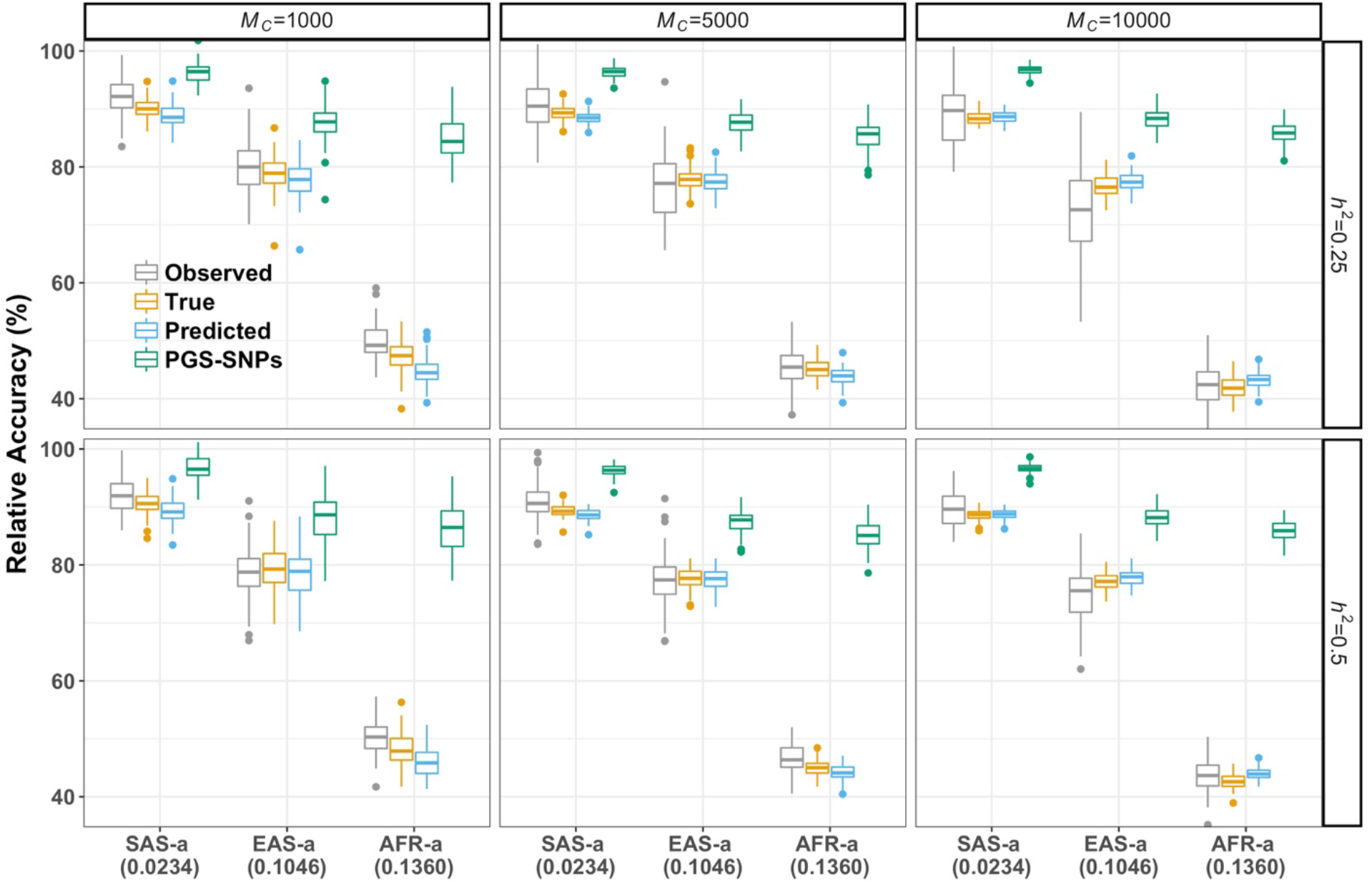
The relative accuracies in different simulation scenarios. We varied the trait heritability (*h*^2^=0.25 and 0.5) and number of *M*_c_ causal variants (*M*_c_=1,000, 5,000 and 10,000) in the simulations. We standardised the accuracies in EUR to 100% and showed the relative accuracies in other target populations using different estimates (see the details in **Material and Methods** section). The predicted accuracy labelled as **“True”** is estimated using Equation (1) based on parameters calculated from SNP pairs of PGS-SNPs and known causal variants within 100 kb; **“Predicted”** accuracy is calculated using SNP pairs of PGS-SNPs and “candidate causal variants” using Equation (2); and **“PGS-SNPs”** is referred to as the estimates using Equation (1) when assuming PGS-SNPs as causal variants. The boxes represent the first and third quantiles and whiskers are 1.5 folds the interquartile range. The points in each box are the estimates in 100 replicates.

### Impact of negative selection

In addition to heritability and polygenicity, another important aspect of the genetic architecture of complex traits and diseases is the relationship between effect sizes at causal variants (hereafter denoted by *β*) and their minor alleles frequencies (hereafter denoted by *p*). This relationship has been modelled in many studies^23–27^ using a parameter *S* such that *β*^2^ is assumed to be proportional to [2*p*(1 − *p*)]^*s*^. Values of *S* determine the relative contributions of common versus rare variants to the genetic variance in the population and thus have been used as an indirect measure of the strength of natural selection^28^. Our model assumes that the variance explained by each causal SNP is constant regardless of allele frequencies. This assumption is consistent with a strong negative selection on causal variants shared between populations and corresponds to a value of *S*=−1. Although previous studies^23,25,29^ have reported pervasive negative selection on complex traits and diseases, these studies often report estimates of *S* with less extreme magnitudes than that assumed in our model. Moreover, given that little is known on the strength of negative selection in non-European populations, we next investigated through additional simulations the impact of violations of this assumption.

We adopted a similar framework (Online Methods) as in our first simulation. However, for the sake of simplicity, we fixed the number of causal variants to *M_c_*=5,000 and the trait heritability to *h*^2^=0.5. We denoted *S*_1_ and *S*_2_ as the value of *S* in Population 1 and Population 2 respectively. We considered three scenarios corresponding to “*S*_1_= *S*_2_=−0.5”, i.e. equal strength of negative selection in both populations, “*S*_1_=−0.75 and *S*_2_=−0.5”, i.e. stronger selection in Population 1 and “*S*_1_=−0.5 and *S*_2_=−0.75”, i.e. stronger selection in Population 2.

In non-European ancestries, we found over 100 replicates that our theory based on Equation (1) predicts a smaller RA than actually observed on average in all three scenarios (**Fig. 2**). This therefore makes our approach conservative. We observed a slightly larger downward bias in the predicted RA when using our heuristic approach based on candidate causal SNPs. On average, the latter approach showed an absolute underestimation of the RA of 1.9% in SAS (i.e. the difference between the observed and the predicted RA is 92.1% −90.1%=1.9%), 6.5% in EAS and 8.5% in AFR, which thus provides a lower bound for the RA. Interestingly, we did not find significant differences in observed and predicted accuracies across scenarios (*t*-test, *t-*statistic=4.69, *p*-value>0.05). This somewhat surprising observation is suggestive that when heritability is constant and effect sizes of causal variants are perfectly correlated, differences in strengths of selection between ancestries might have a negligible impact on the RA of PGS. As a consequence, we can expect differences in strengths of selection between ancestries to mainly impact the term 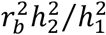 in Equation (1). Note however, that our simulations were based on observed contemporary LD differences between ancestries, which have likely already been shaped by negative selection. This limitation may have masked an additional contribution of differential selection between ancestries.

**Figure 2.**
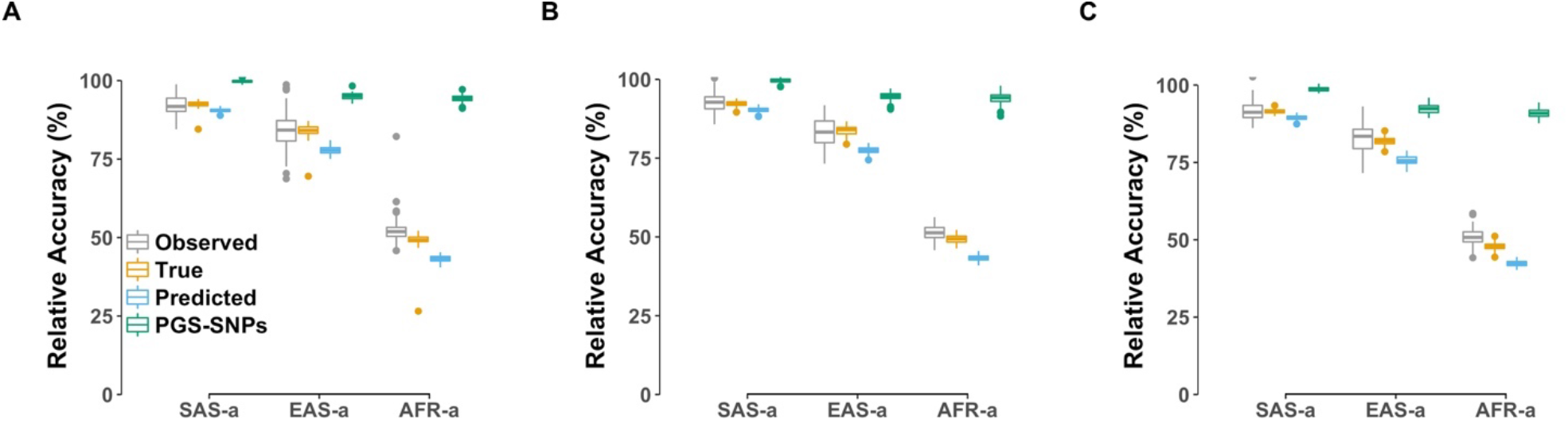
The impact of negative selection on relative accuracies. We showed the relative accuracies in different ancestries under various strengths of negative selection. We simulated the trait with *h*^2^=0.5 and underlain by *M_c_*=5,000 causal variants. We used different strengths of negative selection, modelling as *S* parameter, between discovery and target populations. A) *S*_1_ =*S*_2_ = −0.5; B) *S*_1_ = −0.5, *S*_2_ = −0.75; and C) *S*_1_ = −0.75, *S*_2_ = −0.5. The details of approaches used in results were presented in the **Material and Methods** section, with **“True”** being estimated using Equation (1) based on parameters calculated from SNP pairs of PGS-SNPs and known causal variants within 100 kb; **“Predicted”** accuracies being calculated using SNP pairs of PGS-SNPs and “candidate causal variants” using Equation (2); and **“PGS-SNPs”** referred to as the estimates using Equation (1) when assuming PGS-SNPs as causal variants. The boxes represent the first and third quantiles and whiskers are 1.5 folds the interquartile range. The points in each box are the estimates in 100 replicates.

### Application to real data

We performed GWASs of BMI and height in 313,284 unrelated UKB participants of EUR (**Supplementary Note 4**) and identified 338 and 1,182 quasi-independent GWS SNPs for these two traits respectively (Online Methods). The allele frequency distribution and LD score correlation of those GWS SNPs between ancestries are shown in **Fig. S6** and **Fig. S7**. We used these GWS SNPs to create polygenic predictors of BMI and height and evaluated their predictive performances in the validation and reference sub-samples of the UKB as described in Methods section. Note that we also evaluated the predictive accuracy of PGS based upon sub-significant SNPs (**Supplementary Note 5**) and found, in individuals of non-European ancestries, that PGSs including SNPs selected at less stringent *p*-value thresholds did not systematically improve over using GWS SNPs only (**Fig. 3**). This observation is consistent with previous studies^2,3,18,19^.

**Figure 3.**
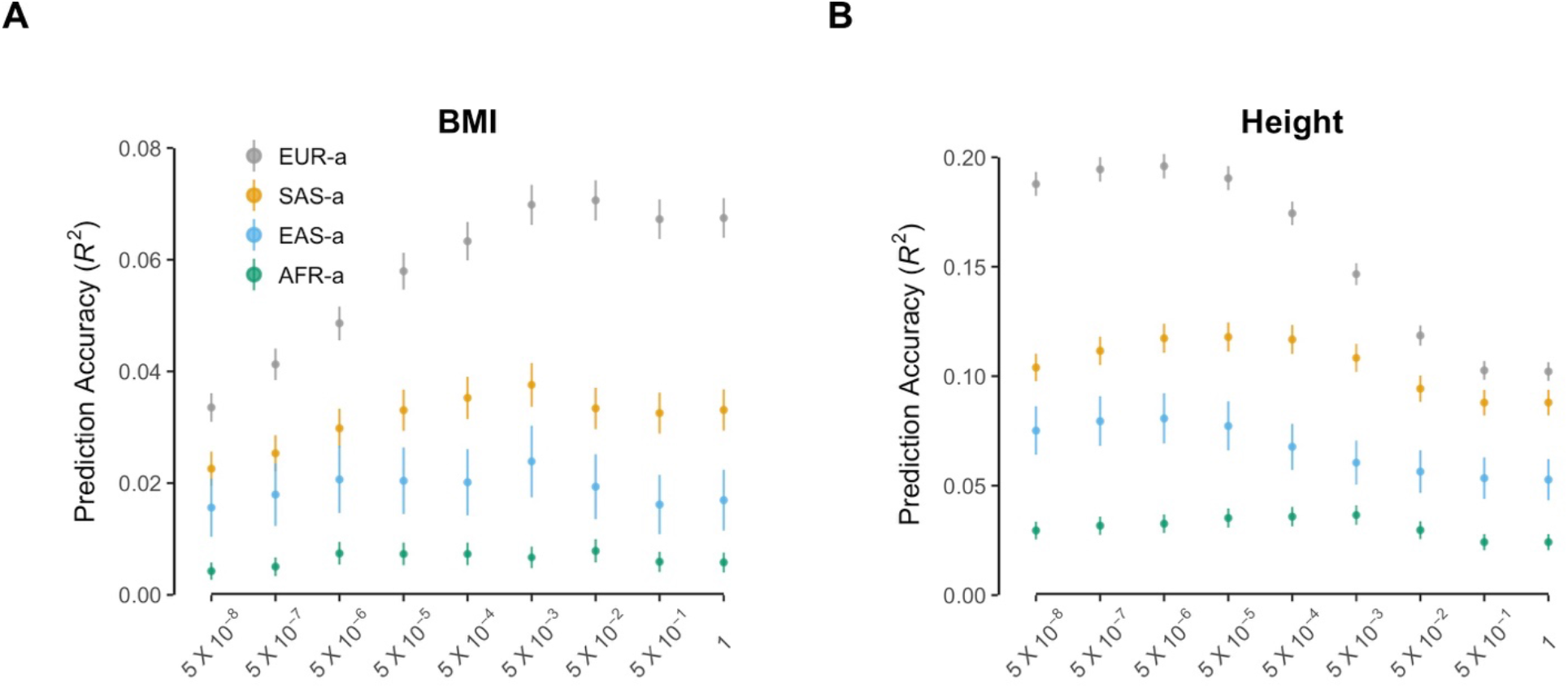
Prediction accuracy at different *p*-value thresholds for BMI and height. We applied a range of *p*-value thresholds to select approximately independent SNPs using the algorithm implemented in LD clumping, details can be seen in **Supplementary Note 5**. The selected SNPs were then used to generate corresponding PGS to estimate the prediction accuracy. Error bars represent the standard errors of the prediction *R*^2^ (see the derivation in **Supplementary Note 6**).

We found that the observed RAs of BMI- and height-PGS based on GWS SNPs reached their minimum values of 12.4% and 15.7% in participants of AFR (**Table 1**). Besides, we observed higher RAs in participants of SAS as compared to those of EAS, which reflects the smaller *F*_ST_ between EUR and SAS (as shown in **Fig. S1**). We then applied our method to predict the accuracy of BMI- and height-PGS (**Fig. 4**) using WGS data from the 1KGP as a reference panel for LD and allele frequency calculations. We define the loss of accuracy (LOA) of PGS as LOA=1-RA and the proportion of LOA explained by LD and MAF as the ratio between the predicted LOA over the observed LOA. Our method predicts that LD and MAF explain 36.6% and 24.3% of the LOA of height- and BMI-PGS in participants of SAS; and up to 70.1% and 71.9% in participants of AFR for BMI and height, respectively. Conversely, these predictions suggest that ~2/3 (~3/4 for height) of the LOA in SAS and ~1/3 of the LOA in AFR is explained by effect sizes heterogeneity (genetic correlation, *r_b_*) and heritability differences between populations^11,30–32^.

**Figure 4.**
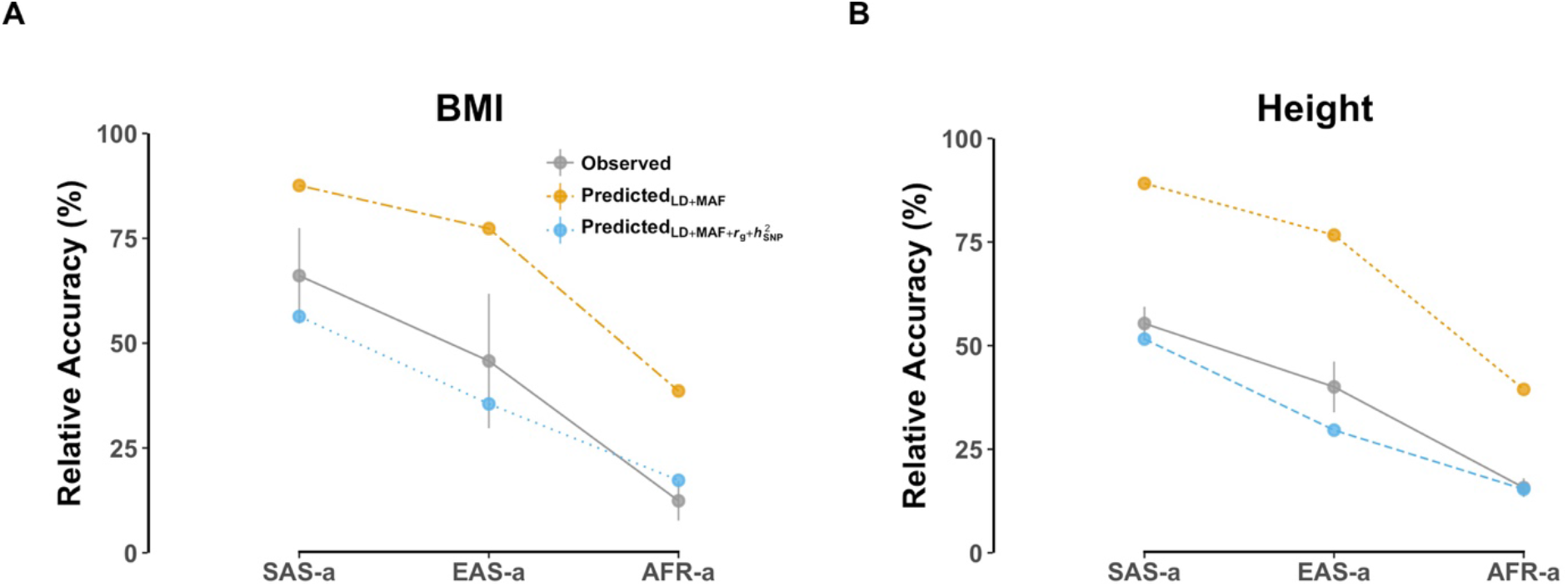
Relative accuracy in different ancestries for BMI and height using GWS SNPs. The **“Observed”** relative accuracies were calculated using approximately independent GWS trait-associated SNPs selected from LD clumping. The error bar represents the standard error for the estimate (see the derivation in **Supplementary Note 6**). We then predicted accuracies deterministically based on GWS SNPs using Equation (2). The results shown as “**Predicted**_LD+MAF_” only used information from LD and MAF differences between ancestries. In addition to LD and MAF differences, we also quantified the contribution of genetic correlation and SNP heritability differences between populations using the Equation (2), which was shown as 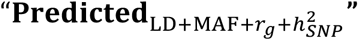 (**Supplementary Note 7**).

**Table 1.** Summary of predictive performance for BMI and height using GWS SNPs.

| **Traits** | **Pops** | **Observed $R^2$** | **Predicted $R^2$** | **Observed RA**<br>**(%)** | **Predicted RA**<br>**(%)** |
| --- | --- | --- | --- | --- | --- |
| **BMI** | **EUR** | 0.034 | 0.034 | 100.0 | 100.0 |
|  | **SAS** | 0.023 | 0.030 | 66.1 | 87.6 |
|  | **EAS** | 0.016 | 0.027 | 45.7 | 77.3 |
|  | **AFR** | 0.004 | 0.013 | 12.4 | 38.6 |
| **Height** | **EUR** | 0.188 | 0.189 | 100.0 | 100.0 |
|  | **SAS** | 0.104 | 0.168 | 55.3 | 89.1 |
|  | **EAS** | 0.075 | 0.145 | 40.0 | 76.7 |
|  | **AFR** | 0.030 | 0.074 | 15.7 | 39.4 |

To further validate our predictions, we approximated the term 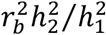 in Equation (2) as 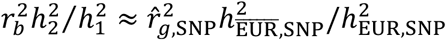, where 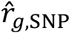 is the joint-SNP effects correlation between ancestries and 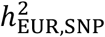 and 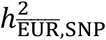 are the SNP heritability in individuals of EUR and non EUR respectively (**Table S1**). We used the bivariate-GREML method implemented in GCTA^33,34^ to estimate 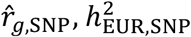 and 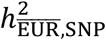 from HapMap3 SNPs (see details in **Supplementary Note 7**). The reliability of this approximation is difficult to assess in practice as it depends on unknown parameters and unknown aspects of the genetic architecture of BMI and height in different ancestries. Nonetheless, we found that predictions including this additional term 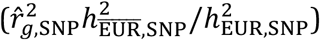 accounted for almost all the remaining LOA of PGS (**Fig. 4**). Yet, we noted that this second approach overall underestimated the RA, which can be expected for at least two reasons. First, our simulation results have shown that if the contribution of rare alleles to heritability is overestimated (i.e. setting *S*=−1 as in our method when in fact *S*>-1) then our method conservatively underestimate the RA of PGS. Secondly, ascertainment of HapMap 3 SNPs, which are more common in EUR, might have led to underestimate 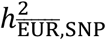 and thus the RA.

Altogether, we have shown here that MAF and LD explain the majority (~70%) of the loss of accuracy of BMI- and height-PGS in individuals of AFR, while the RA of PGS in EAS and SAS is dominated by heritability differences (especially for BMI), thus suggesting a relatively larger contribution of the environment.

## DISCUSSION

In this study, we developed a new theory to predict the relative prediction accuracy of PGS across populations of different ancestry. Our approach only requires GWAS summary statistics as well as data from a globally-diverse reference panel such as the 1KGP. We have shown through simulations that the contribution of differences in LD and MAF to the RA of PGS can be accurately predicted using a simple heuristic approach that models local correlation of LD and MAF across ancestries in the close vicinity of GWS SNPs.

We also explored the impact of negative selection through simulations by considering different relationships between effect sizes and allele frequencies of causal variants. We found that the predicted accuracies were insignificantly different from observed ones, and that they were roughly consistent across different scenarios. However, since PGSs in the current study are mainly focused on common variants, the impact of rarer variants remains to be further investigated.

We further assessed the ability of our theory to predict the accuracy of height- and BMI-PGS in non-European UKB participants. Altogether, we found that up to ~70% of the reduction of RA of height- and BMI-PGS in AFR could be explained by differences in LD and MAF. It is noteworthy that AFR participants of the UK Biobank reside in the UK and therefore are likely to share similar environments as EUR participants. As consequence, the relative contribution of *r_b_* and *h*^2^, which partially reflects the effects of gene by environment interactions (if any), might be underestimated. Similarly, the contribution of 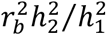 to the RA of PGS in individuals of SAS and EAS might be even larger if evaluated across continents than reported in this study. A recent study by Durvasula et al.^12^ suggested that ~50% of the heritability is captured by European specific variants when a trait is under moderate negative selection, thus limiting the upper bound of prediction accuracy in AFR for example when using European-derived GWAS summary statistics. Our results based on common variants also show a RA<50% in AFR but we demonstrate that this reduction is mostly explained by MAF and LD and not necessarily by population specific causal variants. The latter conclusion is further supported by large genetic correlations (~0.8 on average, **Table S1**) observed between EUR and non-EUR ancestries,^19^ which overall suggests that causal variants underlying common genetic variation identified in European ancestry GWASs are mostly shared across continents.

We note a few limitations to our study. First, our model uses LD and MAF estimated from a reference panel. We have shown the effectiveness of this strategy in population of homogenous ancestry; however, its application remains challenging in admixed populations with complex LD patterns and demographic history. Secondly, we only analyzed common SNPs (with MAF > 0.01) in all populations, which limits the generalization of our conclusions to rarer variants. Indeed, rare variants are more likely to be population specific and are usually poorly imputed using a small imputation reference panel. Lastly, although our simple heuristic strategy to identify “candidate causal variants” worked well in simulations, we expect the use of standard fine-mapping tools to further improve the efficiency of our method.

Despite the acknowledgement of the necessity to collect large scale genome-wide data across different ancestries to fulfil the potential use of PGS in the precision medicine era^3^, this goal remains difficult to achieve in the near future. Instead, trans-ethnic studies have been increasingly popular. They incorporate genotype data from different ethnicities to boost statistic power with increasing sample sizes, which have the benefit to discover disease/trait associated loci and fine-mapping causal variants associated with complex traits or diseases^13,35–38^. However, the structure of the reference population still remains to be thoroughly explored, such as whether some specific populations with certain sample sizes are mostly useful in trans-ethnic studies. Our model presents an opportunity for such study design using both the LD and allele frequency information in a population level. By performing trans-ethnic GWASs, we expect that the predictive ability would increase when the admixed LD structure and allele frequency of the discovery population is similar to the target population.

## ONLINE METHODS

### Samples and quality controls

#### Sample selection

The UK Biobank (**UKB**) comprises of ~500,000 individuals recruited from the UK, aged from 40 to 69 years old. Participants were genotyped using two genotyping arrays, the Affymetrix UK BiLEVE Axiom™ Array and UK Biobank Axiom™ Array. Each participant provided written informed consent. Additional study and quality control details are shown in Bycroft et al.^39^. The approach to infer the ancestry of each individual is described in Yengo et. al^40^. We firstly projected each individual of UKB onto the genotypic principal components (PCs) calculated in 2,000 participants of 1KGP^21^. We only extracted individuals from four ancestries, namely, European ancestry (EUR, N=503), South Asian ancestry (SAS, N=489), East Asian ancestry (EAS, N=504) and African ancestry (AFR, N=504), from 1KGP. We excluded African Caribbeans in Barbados (ACB) and Americans of African Ancestry in SW USA (ASW) populations from AFR and all individuals of American ancestry (AMR) considering their complex admixture patterns. We then assigned each of those genotyped participants of UKB to the closest ancestry based on the first three PCs, resulting in 463,795 EUR, 11,906 SAS, 2,486 EAS and 9,184 AFR. To remove cryptic relatedness in the UKB, we used the GCTA software to calculate the genomic relationship matrix (GRM)^33^ based on genotyped SNPs in each of the aforementioned populations. With one of each pair of individuals with estimated relatedness larger than 0.05 being removed, a subset consisting of unrelated individuals was generated in each ancestry. For the European ancestry, we only extracted those self-reported British and Irish participants. After randomly sampling 10,000 individuals from British subset, we created the discovery dataset using the remaining 313,284 individuals. As for the target populations, we used an independent dataset of ~39,000 UKB individuals. Those individuals included the 10,000 randomly sampled participants who identified themselves as British, 9,979 participants of EUR who identified themselves as Irish, the 9,448 participants of SAS, the 2,257 participants of EAS and the 7,015 participants of AFR. Data from the 1KGP were used as the reference panel in this study. We generated subsets of unrelated individuals in 1KGP with the same strategy as described above, resulting in 495 EUR, 457 SAS, 498 EAS and 484 AFR.

#### SNP selection

For participants of non-European ancestries in the UKB, we further imputed the SNP array data to the 1KGP given that the imputation reference panels, Haplotype Reference Consortium (HRC)^41^ and UK10K^42^, used in the UKB are predominant by European descents. In each ancestry we firstly extracted genotyped SNPs such that Hardy-Weinberg equilibrium (HWE) *p*-value>0.001 and missing rates<0.05 and also excluded individuals with genotype call rates<0.9. Filtered SNPs in each ancestry were then phased using SHAPEIT2^43^ and imputed to 1KGP by IMPUTE2^44^. In each ancestry, stringent quality control procedures were performed separately. We removed SNPs with imputation quality scores<0.30, MAF<0.01, HWE *p*-value<10^-6^, or missing genotype call rates>0.05. HapMap3 SNP set, which has been well designed for human genome-wide common genetic variants^45^, was then extracted from imputed data to run follow-up analyses. A total of 990,395 filtered HapMap3 SNPs in common between populations were selected.

SNPs in the 1KGP reference panel were restricted to a total of 6,877,707 SNPs common to all four ancestries after excluding those SNPs with minor allele count (MAC)<5 in each ancestry. The above described HapMap3 SNP set was further intersected with 1KGP, thus limiting the number to 978,783.

### Simulations

We derived a deterministic equation to quantify the relative prediction accuracy of PGS across ancestries. T o evaluate this equation, we performed various simulations (each scenario with 100 replicates) using the real genotypes in the UKB cohort. To simplify our notations, we denoted the training/discovery sample as Population 1 (*l* = 1) and the target/test sample as Population 2 (*l* = 2).

The phenotypes were simulated based on the additive model *y* = *g* + *e* in all scenarios using different values of *M_c_* and trait heritability (*h*^2^). In each simulation replicate, we simulated a trait with a heritability of 0.25 or 0.5 in all ancestries. Traits were simulated from *M_c_*(*M_c_*=1,000, 5,000, and 10,000) causal variants sampled at random from the HapMap3 SNPs. We assumed the effect sizes of causal variants (*β*) were perfectly correlated across populations, i.e. *r_b_* = 1. For each causal variant, *β* was sampled from a normal distribution with mean 0 and variance 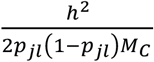, where *p_jl_* is the MAF in *j*^th^ causal variant in population *l.* For each individual, the genetic value *g* was defined such as 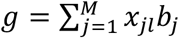, where *x_jl_* denotes the minor allele count (*x_jl_* equals to 0, 1 or 2) at the *j*^th^ causal variant in population *l*. The environmental effect (*e*) was simulated using a normal distribution with mean 0 and variance equal to (1 − *h*^2^): *e* ~ *N*(0,1 − *h*^2^), such that the phenotypic variance across populations was equal to 1.

PLINK1.90 (note version 20 Mar was used in this study)^46^ was used to run GWAS for the simulated phenotypes in Population 1 using the simple linear association testing on HapMap3 SNPs. To mimic the imperfect LD between the causal variants and the SNP markers used in GWAS, the causal variants were always left out of the analysis.

To further explore the impact of negative selection, we sampled *β* from a multi-normal distribution with mean 0 and variance 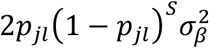, where 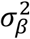 is the variance of causal effect sizes. We considered three scenarios corresponding to “*S*_1_=*S*_2_=−0.5”, i.e. equal strength of selection in both ancestries, “*S*_1_=−0.75 and *S*_2_=−0.5”, i.e. stronger selection in Population 1 and “*S*_1_=−0.5 and *S*_2_=−0.75”, i.e. stronger selection in Population 2. The phenotypes were generated in the same way as described above. For simplicity, we focused on a trait with a heritability *h*^2^=0.5 and controlled by *M_c_*=5,000 causal variants.

### Selection of genome-wide significant (GWS) SNPs

After running GWAS, we selected approximately independent SNPs associated with the trait (referred to as “GWS SNPs” or “PGS-SNPs”), using the “LD clumping” algorithm implemented in PLINK1.90^46^. We used the following command: --*clump-p1 5e-8 --clump-p2 5e-8 --clump-kb 2000 --clump-r2 0.01.* The genotypes of the training population were used as LD reference for clumping. We used here a more stringent LD threshold than classically used (e.g. 0.1 or 0.2) because SNPs with LD *r*^2^ as large as 0.1 can still reflect the same signal when GWAS sample size is large (e.g. N>300,000).

### Deterministic accuracies of genetic prediction

We calculated the deterministic accuracy in the target populations based on selected GWS SNPs. Since we focused on in this study the contribution of LD and allele frequency differences between populations to the RA, the term 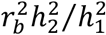 equalled to 1 in simulations. We then used a reference panel to calculate the LD correlation and MAF between populations. To first validate our theory, we used Equation (1), which assumes causal variants to be known, to calculate the LD correlation and MAF using SNP pairs between PGS-SNPs and “known causal variants”. We then explored the performance of our heuristic method using Equation (2), given that causal variants are typically unknown or unobserved. For that, we took advantage of the LD and MAF information between PGS-SNPs and “candidate causal variants” instead. Further, we applied a naïve approach assuming PGS-SNPs to be the causal variants, thus mainly the allele frequency differences between populations would be captured using Equation (1).

When using Equation (1) with known causal variants, we firstly matched GWS SNPs to them to calculate LD correlation and allele frequencies between populations (results shown as **“True”** in all figures). It was done by constraining the window centred at each GWS SNP as 100 kb and then selecting those pairs including known causal variants. This window was based on the report that ~95% top lead SNPs (with MAF > 0.01) identified from GWASs are within 100 kb distance from the causal variants in European ancestry^20^. Although the causal variants are often unknown or unobserved in a classical GWAS, they are usually tagged by numerous SNPs. Therefore, we took advantage of the information regarding fine-mapping precision of GWAS studies and selected “candidate causal variants” as those SNPs in LD *r*^2^>0.45 with GWS SNPs and located within 100 kb window^20^. Those GWS SNPs and “candidate causal variants” pairs were then used in Equation (2), with results referring to as **“Predicted”** in all figures. When assuming the PGS-SNPs as causal variants, we estimated the accuracies using Equation (1) where the LD correlation was replaces with 1 (results denoted as **“PGS-SNPs”** in all figures).

The final predicted parameters to evaluate predictive ability were calculated as the mean of the estimates across 100 replicates, respectively. To explore the impact of imputation on our model, we used both 1KGP WGS data and UKB imputed data as reference in the simulated data. We used the same approaches, except for the one assuming causal variants were known, to analyse GWAS summary statistics of BMI and height (**Supplementary Note 4**). Whilst, 1KGP WGS data were used as the reference panel since most causal variants might not be included in the summary statistics. We further explored the impact of the term 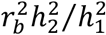 on the predictive performance by estimating the genetic correlation and SNP heritability between ancestry (**Supplementary Note 7**).

The relative accuracy (RA) was calculated as the ratio of 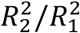, where 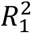 was the predicted prediction accuracy in the population with same ancestry of discovery population; and 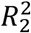 was the predicted prediction accuracy in other target populations. Note we have two target populations of EUR, so we used each of them as the validation, in which the accuracy of PGS within-European-ancestry was quantified and a reference group, in which we predicted the accuracy of PGS, separately. We then averaged the accuracies for each target populations.

### Empirical accuracy of genetic prediction

After selecting approximately independent GWS SNPs using LD clumping, we then generated PGS in each target population by adding up the product of minor allele counts times effect sizes of GWS SNPs estimated from GWAS summary statistics. The prediction accuracy was then estimated using the squared correlation (*R*^2^) between the true phenotypes and the PGS. The empirical relative accuracies were calculated in a similar way as deterministic accuracies mentioned above.

## Acknowledgements

This research was supported by the The Australian National Health and Medical Research Council (1113400) and the Australian Research Council (DP160102400, FT180100186 and FL180100072). The funders had no role in study design, data collection and analysis, decision to publish, or preparation of the manuscript. This research has been conducted using the UK Biobank Resource under project 12505. We thank all participants of the UK Biobank.

## Author Contributions

LY and PMV conceived the study. LY derived the theory. LY and YW designed the experiment. YW prepared data and performed analyses under the assistance and guidance from JG, GN, JY, PMV and LY. YW and LY wrote the manuscript with the participation of all authors. All authors reviewed and approved the final manuscript.

## URLs

UKB consortium: http://www.ukbiobank.ac.uk/

GCTA: http://cnsgenomics.com/software/gcta

PLINK: https://www.cog-genomics.org/plink2

IMPUTE2: https://mathgen.stats.ox.ac.uk/impute/impute_v2.html

SHAPEIT2: https://mathgen.stats.ox.ac.uk/genetics_software/shapeit/shapeit.html

1KGP: ftp://ftp.1000genomes.ebi.ac.uk/vol1/ftp/data_collections/1000_genomes_project/data

## Data availability

This study makes use of genotype and phenotype data from the UK Biobank data under project 12505. UKB data can be accessed upon request once a research project has been submitted and approved by the UKB committee.

