## Supplementary Notes for "Theoretical and empirical quantification of the accuracy of polygenic scores in ancestry divergent populations"

##### 1. Theoretical expectation of trans-ethnic accuracy of polygenic scores

We assume causal variants to be shared between ancestries but allow their effect sizes to vary from one ancestry to another. Therefore, ancestry-specific causal variants are a special case with non-zero effect sizes in only one ancestry. We also assume that effect sizes are correlated between population and denote $r_b$ that correlation. We finally assume that the trait is controlled by $M_C$ causal variants and that prediction is based on $M_T$ independent genome-wide significant (GWS) SNPs.

We can write the phenotypic value of individual from population $l$ as

(1) $$y_l = \sum_{j=1}^{M_C} b_{jl}\left(\frac{x_{jl}^{(c)} - 2p_{jl}^{(c)}}{\sqrt{2p_{jl}^{(c)}\left(1-p_{jl}^{(c)}\right)}}\right) + e_l,$$

where $x_{jl}^{(c)}$ is the minor allele count (MAC) at the $j^{\text{th}}$ causal variant in population $l$, $b_{jl}$ is the effect size per standardized genotype at the $j^{\text{th}}$ causal variant in population $l$, $p_{jl}^{(c)}$ is the minor allele frequency (MAF) at the $j^{\text{th}}$ causal variant in population $l$, and $e_l$ is a residual term measuring non-genetic effects in population $l$. We hereafter denote $h_{jl}^{(c)} = 2p_{jl}^{(c)}\left(1-p_{jl}^{(c)}\right)$.

We now consider that the discovery GWAS is performed in individuals of Population 1 and accuracy is evaluated in individuals of Population 2. We denote $\hat{y}_l$ as the PGS calculated in individuals of population $l$. The squared correlation $R_2^2$ between $y_2$ and $\hat{y}_2$ can be expressed as

(2) $$R_2^2 = \frac{\text{cov}^2(y_2,\hat{y}_2)}{\text{var}(\hat{y}_2)\text{var}(y_2)} = \left[\frac{\text{cov}(y_2,\hat{y}_2)}{\text{cov}(y_1,\hat{y}_1)}\right]^2 \times \frac{\text{cov}^2(y_1,\hat{y}_1)}{\text{var}(\hat{y}_1)\text{var}(y_1)} \times \frac{\text{var}(\hat{y}_1)}{\text{var}(\hat{y}_2)} \times \frac{\text{var}(y_1)}{\text{var}(y_2)}.$$

We assume, without loss of generality, that $\text{var}(y_1) = \text{var}(y_2) = 1$. Therefore, Equation (2) can be rewritten as

(3) $$R_2^2 = R_1^2 \times \left[\frac{\text{cov}(y_2,\hat{y}_2)}{\text{cov}(y_1,\hat{y}_1)}\right]^2 \times \frac{\text{var}(\hat{y}_1)}{\text{var}(\hat{y}_2)},$$

where $R_1^2$ denotes the prediction accuracy in individuals of same ancestry as in the discovery sample, i.e. Population 1.

Equation (3) depends on two parameters, which can be evaluated without observing the phenotype in the target population. Those parameters are $R_1^2$ which can be obtained from previous prediction studies in individuals of Population 1; and $\mathrm{var}(\hat{y}_1)/\mathrm{var}(\hat{y}_2)$, which can be evaluated using data from a reference panel. The only unspecified parameter is $\mathrm{cov}(y_2,\hat{y}_2)/\mathrm{cov}(y_1,\hat{y}_1)$ for which we derive below an approximation.

We assume causal SNP effect sizes to be normally distributed, i.e.

(4) $$\begin{bmatrix} b_{j1} \\ b_{j2} \end{bmatrix} \sim \mathcal{N}\left(\begin{bmatrix} 0 \\ 0 \end{bmatrix}, \ \frac{1}{M_C}\begin{bmatrix} h_1^2 & r_b\sqrt{h_1^2 h_2^2} \\ r_b\sqrt{h_1^2 h_2^2} & h_2^2 \end{bmatrix}\right),$$

where $h_l^2$ denotes the heritability in population $l$.

Conditional on the $b_{j1}$'s, the ordinary least squared (OLS) estimator $\hat{\beta}_{\mathrm{k}1}^{(\mathrm{t})}$ of SNP $k$ calculated in individuals of Population 1 is such that

(5) $$\mathrm{E}[\hat{\beta}_{k1}^{(t)}] \ = \ \frac{\mathrm{cov}(x_{k1},y_1)}{\mathrm{var}(x_{k1})} = \sum_{j=1}^{M_C} \frac{b_{j1}}{\sqrt{h_{j1}^{(c)}}} \times \frac{\mathrm{cov}(x_{k1}^{(t)},x_{j1}^{(c)})}{\mathrm{var}(x_{k1}^{(t)})} = \frac{1}{\sqrt{h_{k1}^{(t)}}} \sum_{j=1}^{M_C} b_{j1} r_{jk,1} \,,$$

where $h_{k1}^{(t)} = var(x_{k1}) = 2p_{j1}^{(t)}\left(1-p_{j1}^{(t)}\right)$ and $r_{jk,\mathrm{l}} = cov\left(x_{k1}^{(t)}, x_{j1}^{(c)}\right)/\sqrt{h_{j1}^{(c)} \times h_{k1}^{(t)}}$ is the LD correlation between causal SNP $j$ and GWS SNP $k$ in population $l$.

We can also express the variance of $\hat{\beta}_{\mathrm{k}1}^{(\mathrm{t})}$ as

(6) $$\mathrm{var}(\hat{\beta}_{k1}^{(t)}) = \frac{\mathrm{var}(y_1) - \mathrm{E}[\hat{\beta}_{k1}^{(t)}]^2 \mathrm{var}(x_{k1}^{(t)})}{N_1 \mathrm{var}(x_{k1}^{(t)})} = \frac{\mathrm{var}(y_1) - \mathrm{E}[\hat{\beta}_{k1}^{(t)}]^2 h_{k1}^{(t)}}{N_1 h_{k1}^{(t)}} = \frac{\mathrm{var}(y_1)}{N_1 h_{k1}^{(t)}} - \frac{1}{N_1}\mathrm{E}[\hat{\beta}_{k1}^{(t)}]^2.$$

In the target sample of Population 2, $\hat{y}_2$ is defined as

(7) $$\hat{y}_2 = \sum_{k=1}^{M_T} \hat{\beta}_{k1}^{(t)} x_{k2}^{(t)} = \mu_{\mathrm{PGS}} + \sum_{k=1}^{M_T} \hat{\beta}_{k1}^{(t)}\left[x_{k2}^{(t)} - 2p_{k2}^{(t)}\right],$$

where $\mu_{\mathrm{PGS}} = \sum_{k=1}^{M_T} \hat{\beta}_{k1}^{(t)} 2p_{k2}^{(t)}$.

We note that $\mu_{\mathrm{PGS}}$ is constant for all individuals in the target sample. Therefore, conditional on SNP true effect sizes $\boldsymbol{b} = \left((b_{11},b_{12}),\ldots,(b_{j1},b_{i2}),\ldots,(b_{M_C1},b_{M_C2})\right)$, the

(8) $$\mathrm{cov}(\hat{y}_2,y_2 \,|\, \boldsymbol{b}) \ = \mathrm{cov}\left(\sum_{k=1}^{M_T} \hat{\beta}_{k1}^{(t)}\left[x_{k2}^{(t)} - 2p_{k2}^{(t)}\right], y_2\right).$$

$$= \text{cov}\left[\sum_{k=1}^{M_T} \hat{\beta}_{k1}^{(t)}[x_{k2}^{(t)} - 2p_{k2}^{(t)}], \sum_{j=1}^{M_C} b_{j2}\left(\frac{x_{j2}^{(c)} - 2p_{j2}^{(c)}}{\sqrt{h_{j2}^{(c)}}}\right) + e_2\right]$$

$$= \sum_{k=1}^{M_T}\sum_{j=1}^{M_C} \mathrm{E}[\hat{\beta}_{k1}^{(t)}] b_{j2}\sqrt{h_{k2}^{(t)}} \times \frac{\text{cov}(x_{k2}^{(t)}, x_{j2}^{(c)})}{\sqrt{h_{j2}^{(c)} h_{k2}^{(t)}}}$$

$$= \sum_{k=1}^{M_T}\sum_{j=1}^{M_C} \sqrt{\frac{h_{k2}^{(t)}}{h_{k1}^{(t)}}} \times b_{j2} r_{jk,2}\left(\sum_{i=1}^{M_C} b_{i1} r_{ik,1}\right)$$

$$= \sum_{k=1}^{M_T} \sqrt{\frac{h_{k2}^{(t)}}{h_{k1}^{(t)}}}\left(\sum_{j=1}^{M_C} b_{j1} b_{j2} r_{jk,1} r_{jk,2} + \sum_{i \neq j} b_{i1} b_{j2} r_{ik,1} r_{jk,2}\right)$$

Similarly, we can show that

(9) $\quad \text{cov}(\hat{y}_1, y_1 | \boldsymbol{b}) = \sum_{k=1}^{M_T}\left(\sum_{j=1}^{M_C} b_{j1}^2 r_{jk,1}^2 + \sum_{i \neq j} b_{i1} b_{j2} r_{ik,1} r_{jk,2}\right).$

We finally, approximate $\text{cov}(y_2, \hat{y}_2)/\text{cov}(y_1, \hat{y}_1)$ with its expectation as

(10) $\quad \frac{\text{cov}(y_2,\hat{y}_2)}{\text{cov}(y_1,\hat{y}_1)} \approx \mathbb{E}\left(\frac{\text{cov}(y_2,\hat{y}_2)}{\text{cov}(y_1,\hat{y}_1)}\right) \approx \frac{\mathbb{E}(\mathbb{E}[\text{cov}(\hat{y}_2, y_2|\boldsymbol{b})])}{\mathbb{E}(\mathbb{E}[\text{cov}(\hat{y}_1, y_1|\boldsymbol{b})])}$

Under the assumption that effect sizes of causal SNPs are independent random variables, we have that $\mathbb{E}[b_{il} b_{jl}] = 0$ when $i \neq j$, and given that $\mathbb{E}[b_{j1} b_{j2}] = r_b\sqrt{h_1^2 h_2^2}/M_C$ we can write

(11) $\quad \frac{\text{cov}(y_2,\hat{y}_2)}{\text{cov}(y_1,\hat{y}_1)} \approx r_b \times \sqrt{\frac{h_2^2}{h_1^2}} \times \frac{\sum_{k=1}^{M_T}\sqrt{\frac{h_{k2}^{(t)}}{h_{k1}^{(t)}}}\left(\sum_{j=1}^{M_C} r_{jk,1} r_{jk,2}\right)}{\sum_{k=1}^{M_T}\left(\sum_{j=1}^{M_C} r_{jk,1}^2\right)},$

hence Equation (1) in the main text.

**2. Wright's $F_{ST}$ calculation**

The pairwise Wright's $F_{ST}$[1] between discovery and target dataset was used to estimate the genetic distance. For the sake of computational efficiency, we randomly sampled 50,000 individuals from the discovery dataset as the reference and calculated the $F_{ST}$ using the software PLINK1.90[2] based on HapMap3 SNPs.

**3. Estimates of allele frequency and LD score between ancestries**

To show the allele frequency and LD differences of GWS SNPs for height and BMI between ancestries, we calculated the allele frequencies and LD scores using 1KGP WGS data as the reference panel. The LD score in this study was calculated as 1 + $\overline{r^2} \times m$, where $\overline{r^2}$ is the mean LD $r^2$ between the target SNP and all other SNPs in the window and $m$ is the number of SNPs in the window. In each ancestry, we calculated the LD score within 100 kb window using the software GCTA[3]. The 100 kb window was consistent with the arbitrary choice we made in the estimation of LD correlation in our theory. We used the frequency based on the same allele to estimate the allele frequency correlation between ancestries.

To further explore the impact of imputation on the estimates of LD correlation and MAF in our theory, we also calculated the LD scores and MAF based on HapMap3 SNPs using UKB imputed data. We used the heterozygosity calculated as $2p_{ij}(1-p_{ij})$, with $p$ being the MAF of the $i^{\text{th}}$ SNPs in the $j^{\text{th}}$ population, to estimate the allele frequency correlation between ancestries.

#### 4. GWAS for height and BMI in UK Biobank

The phenotypes of height and BMI were pre-adjusted by age, sex, recruitment centre, genotyping batches and the first 10 PCs. The residuals of them were then inverse-normal transformed with the aim to make them more normally distributed. Simple linear association tests were performed in 313,284 British as described in **Material and Methods** using PLINK1.90[2].

#### 5. Predictive performance using sub-significant SNPs

We also evaluated the predictive performance of PGS for height and BMI using sub-significant SNPs. We used a range of $p$-value thresholds to select PGS SNPs, i.e. $5 \times 10^{-7}$, $5 \times 10^{-6}$, $5 \times 10^{-5}$, $5 \times 10^{-4}$, $5 \times 10^{-3}$, $5 \times 10^{-2}$, $5 \times 10^{-1}$ and 1. The process was the same as described in **Material and Methods** to select the GWS SNPs. After selecting the PGS SNPs, we generated corresponding PGS and then calculated the prediction $R^2$ (see **Material and Methods**) in the target populations.

#### 6. Standard errors for the observed relative accuracies

We derive standard errors for the observed relative accuracies $\hat{R}_2^2/\hat{R}_1^2$ using the Delta-method based on Taylor series expansions. We first recall the asymptotic variance of the Pearson correlation coefficient $\hat{r}$, estimated in sample of $N$ individuals:

(12) $\text{var}(\hat{r}) \approx \frac{1-\hat{r}^2}{N}$.

Then, using the Delta-method, we can show that

(13) $\mathrm{var}(\hat{r}^2) \approx 4\hat{r}^2(1-\hat{r}^2)/N$.

Noting that the squared derivative of the function $f(x) = x^2$, is $[f'(x)]^2 = 4x^2$. Moreover, the variance of ratios lemma[4] posits that

(14) $\mathrm{var}(u/v) \approx \left[\frac{\mathbb{E}(u)}{\mathbb{E}(v)}\right]^2 \left(\frac{\mathrm{var}(u)}{[\mathbb{E}^2(u)]} - \frac{2\mathrm{cov}(u,v)}{[\mathbb{E}(u)\mathbb{E}(v)]} + \frac{\mathrm{var}(v)}{[\mathbb{E}^2(v)]}\right)$

Therefore, assuming $\mathbb{E}(\hat{u}) \approx \hat{u}$, and given that estimated accuracies $\hat{R}_1^2$ and $\hat{R}_2^2$ are independent (i.e. $\mathrm{cov}\left(\hat{R}_1^2, \hat{R}_2^2\right) = 0$), we can finally write

(15) $\mathrm{var}(\hat{R}_2^2/\hat{R}_1^2) \approx \left[\frac{\hat{R}_2^2}{\hat{R}_1^2}\right]^2 \left(\frac{4(1-\hat{R}_1^2)}{N_1\hat{R}_1^2} + \frac{4(1-\hat{R}_2^2)}{N_2\hat{R}_2^2}\right)$

**7. Impact of genetic correlation and SNP heritability difference in real data**

To explore the impact of genetic correlation ($r_b$ in this study) and SNP heritability difference ($h_2^2/h_1^2$) on the RA (see Equation (2) in the main text), we performed bi-variate GREML analysis implemented in GCTA[3,5] to estimate the joint-SNP effects correlation ($\hat{r}_{g,\mathrm{SNP}}$) and SNP heritability ($h_{SNP}^2$) for height and BMI, respectively. The analysis was performed using HapMap3. For the time efficiency, we used the aforementioned subset of British individuals (N=50,000) as a reference for the discovery population. The genotypes between the discovery and each of the target populations were combined; and we used the *"--sub-popu"* option in GCTA to build the GRM based on population specific allele frequency. We then approximated $r_b^2 h_2^2/h_1^2$ as $\hat{r}_{g,\mathrm{SNP}}^2 h_{\overline{\mathrm{EUR}},\mathrm{SNP}}^2/h_{\mathrm{EUR},\mathrm{SNP}}^2$ ($h_{\mathrm{EUR},\mathrm{SNP}}^2$ and $h_{\overline{\mathrm{EUR}},\mathrm{SNP}}^2$ are the SNP heritability in individuals of EUR-a and non EUR-a respectively) and re-predicted the accuracy using Equation (2).

### Supplementary Figures and Tables

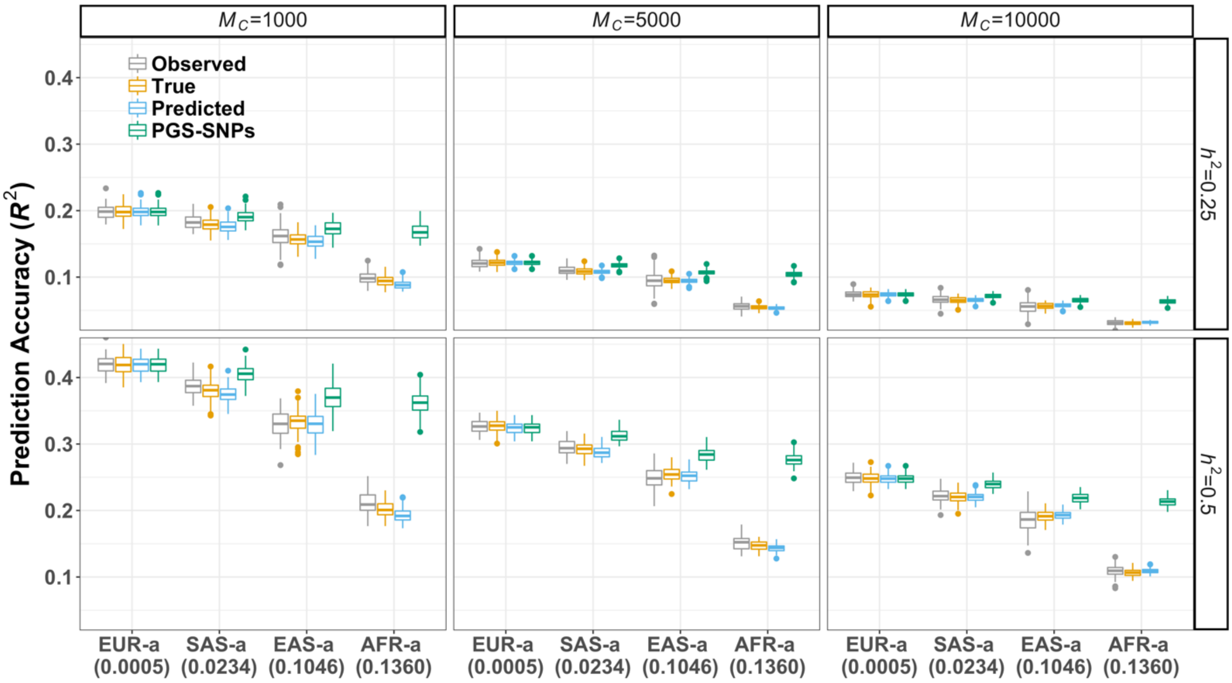


**Figure S1. Predictive performance in different simulation scenarios.** We varied the trait heritability ($h^2$=0.25 and 0.5) and number of $M_C$ causal variants ($M_C$=1,000, 5,000 and 10,000). The approaches to estimate different accuracies were detailed in **Material and Methods** section. The **"Observed"** is the observed prediction $R^2$ in different ancestries. The predicted accuracy labelled as **"True"** is estimated using Equation (1) based on parameters calculated from SNP pairs of PGS-SNPs and known causal variants within 100 kb; **"Predicted"** accuracy is calculated using SNP pairs of PGS-SNPs and "candidate causal variants" using Equation (2); and **"PGS-SNPs"** is referred to as the estimates using Equation (1) when assuming PGS-SNPs as causal variants. The numbers under the ancestry labels in x-axis denoted the pairwise $F_{ST}$ calculated using HapMap 3 SNPs between discovery population and target population (see **Supplementary Note 2**). The boxes represent the first and third quantiles and whiskers are 1.5 folds the interquartile range. The points in each box are the estimates in 100 replicates.

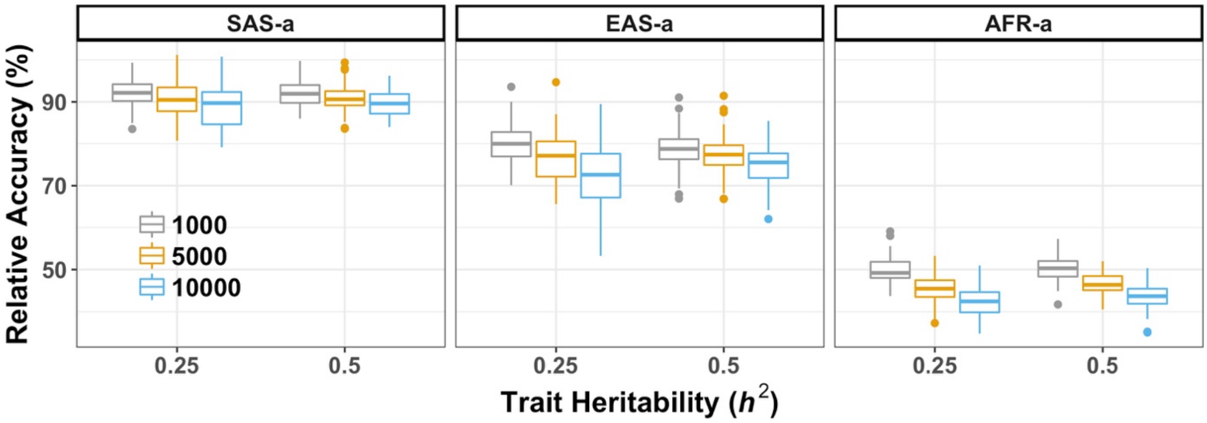


**Figure S2. The observed relative accuracies in different simulation scenarios.** We varied the trait heritability ($h^2$=0.25 and 05) and number of $M_C$ causal variants ($M_C$=1,000, 5,000 and 10,000) in the simulations. As for the relative accuracies, we showed they were relatively consistent across scenarios in different ancestries, although a slight downward trend was presented with the increasing number of $M_C$. The boxes represent the first and third quantiles and whiskers are 1.5 folds the interquartile range. The points in each box are the estimates in 100 replicates.

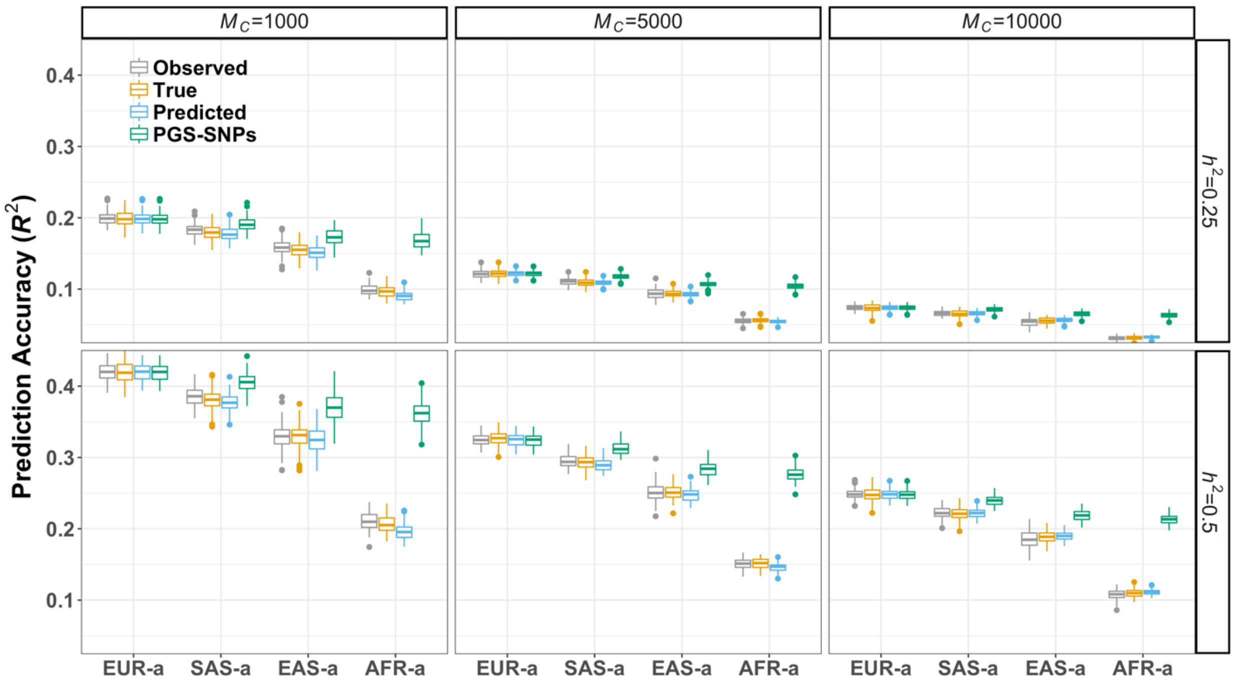


**Figure S3. Predictive performance in different simulation scenarios using UKB imputed data as the reference panel.** We varied the trait heritability ($h^2$=0.25 and 0.5) and number of $M_C$ causal variants ($M_C$=1,000, 5,000 and 10,000) in the simulations. The **"Observed"** is the prediction $R^2$ in different ancestries. The predicted accuracies labelled as **"True"** were estimated using Equation (1) based on parameters calculated from SNP pairs of PGS-SNPs and known causal variants within 100 kb; **"Predicted"** accuracies were calculated using SNP pairs of PGS-SNPs and "candidate causal variants" using Equation (2); and **"PGS-SNPs"** referred to as the estimates using Equation (1) when assuming PGS-SNPs as causal variants. We showed the results across scenarios were quite consistent with those obtained using 1KGP WGS data as the reference (**Fig. S1**). The boxes represent the first and third quantiles and whiskers are 1.5 folds the interquartile range. The points in each box are the estimates in 100 replicates.

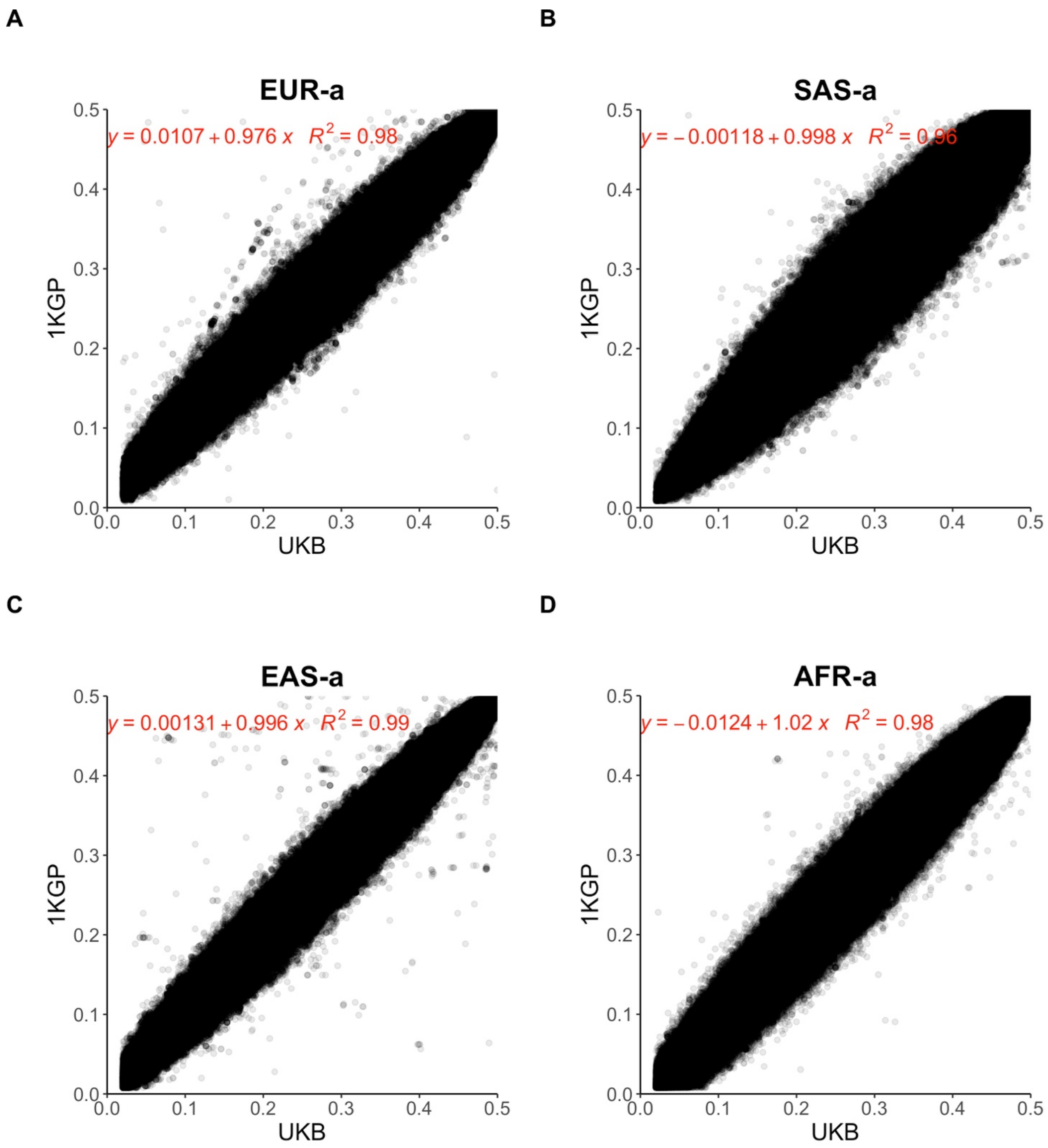


**Figure S4. The allele frequency correlation between populations using different reference panels.** We calculated the SNP heterozygosity as $2p(1-p)$ in each ancestry from 1KGP WGS and UKB imputed data, respectively (see details in **Supplementary Note 3**). We found the results was quite consistent between these two datasets.

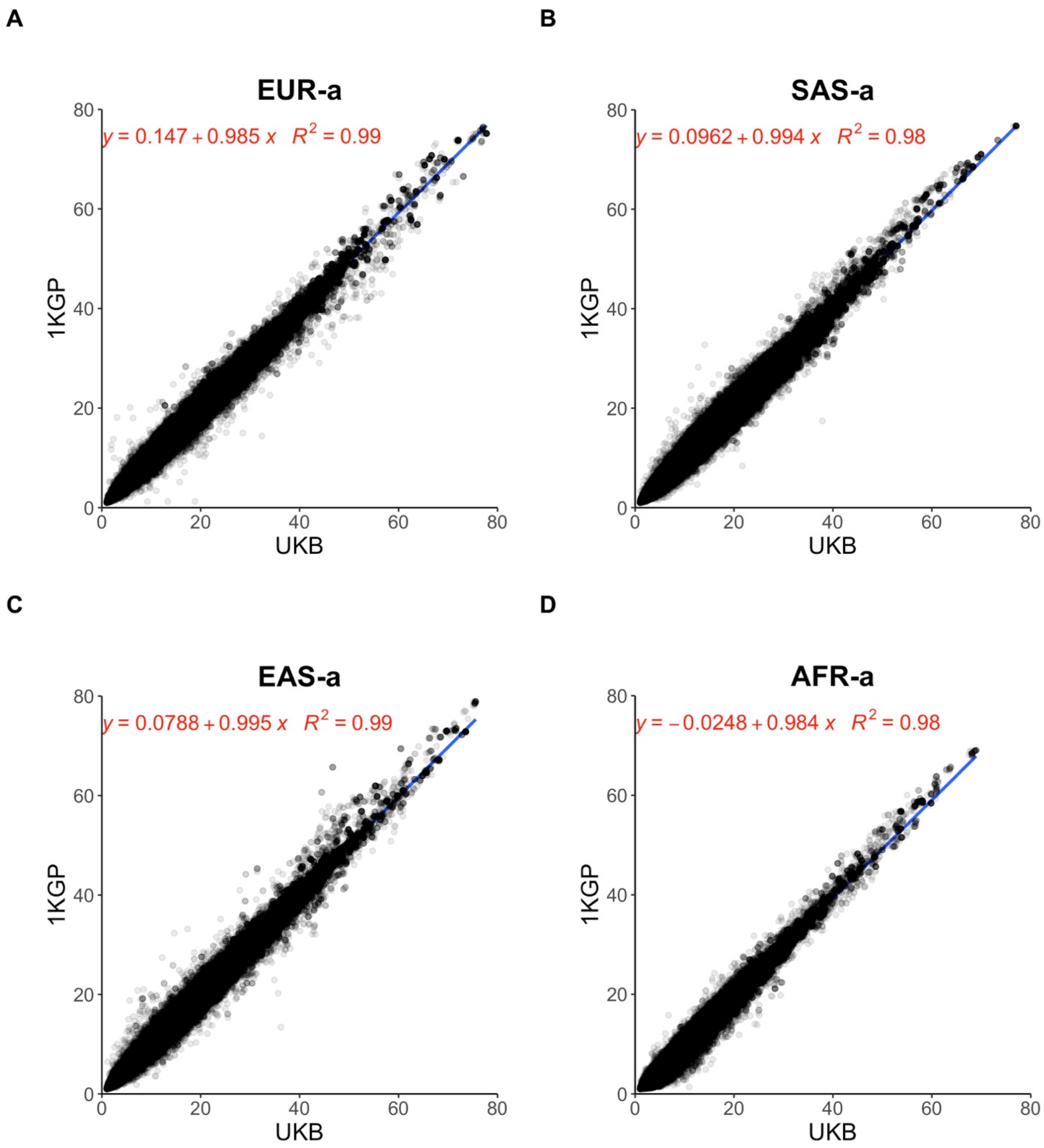


**Figure S5. The LD score correlation between populations using different reference panels.** We calculated the LD score of HapMap3 SNPs within 100 kb window in each ancestry from 1KGP WGS and UKB imputed data, respectively (see details in **Supplementary Note 3**). We found the LD score was quite consistent either using 1KGP WGS or UKB imputed data as the reference panel.

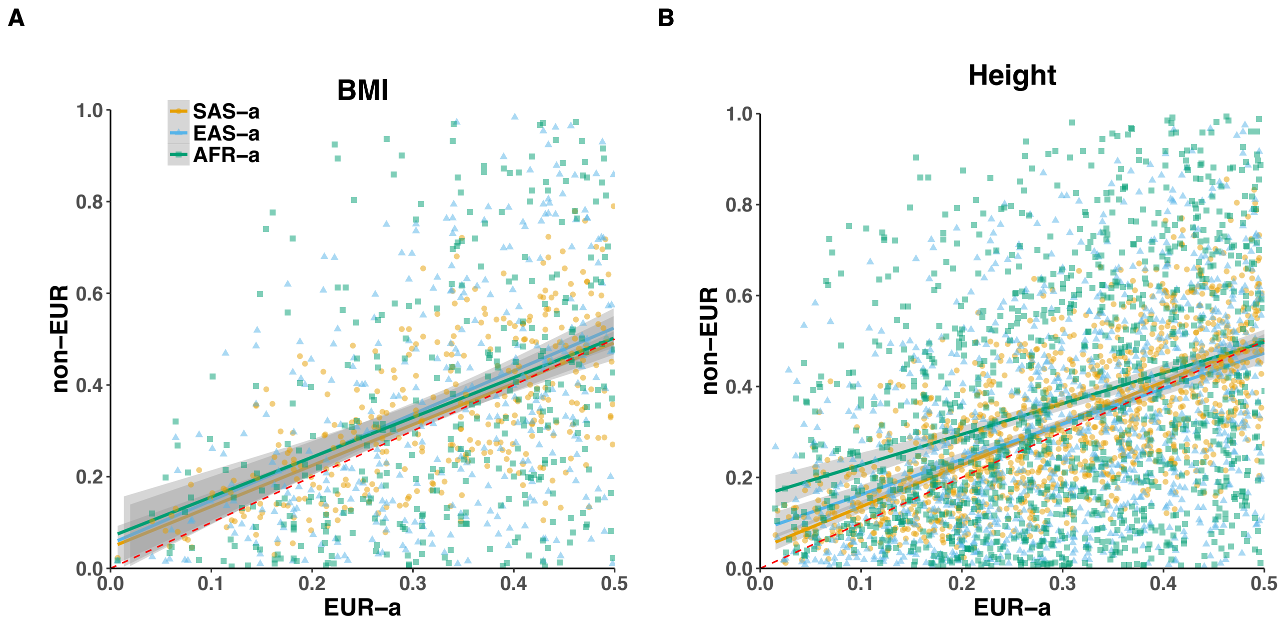


**Figure S6. The allele frequency distribution of trait-associated loci between ancestries.** We compared the allele frequency distributions of the GWS SNPs between the discovery population (EUR-a) and other non-European target populations (non-EUR). The comparison was done using the same allele of those GWS SNPs for height and BMI, respectively. The red dashed line in each plot was: y=x.

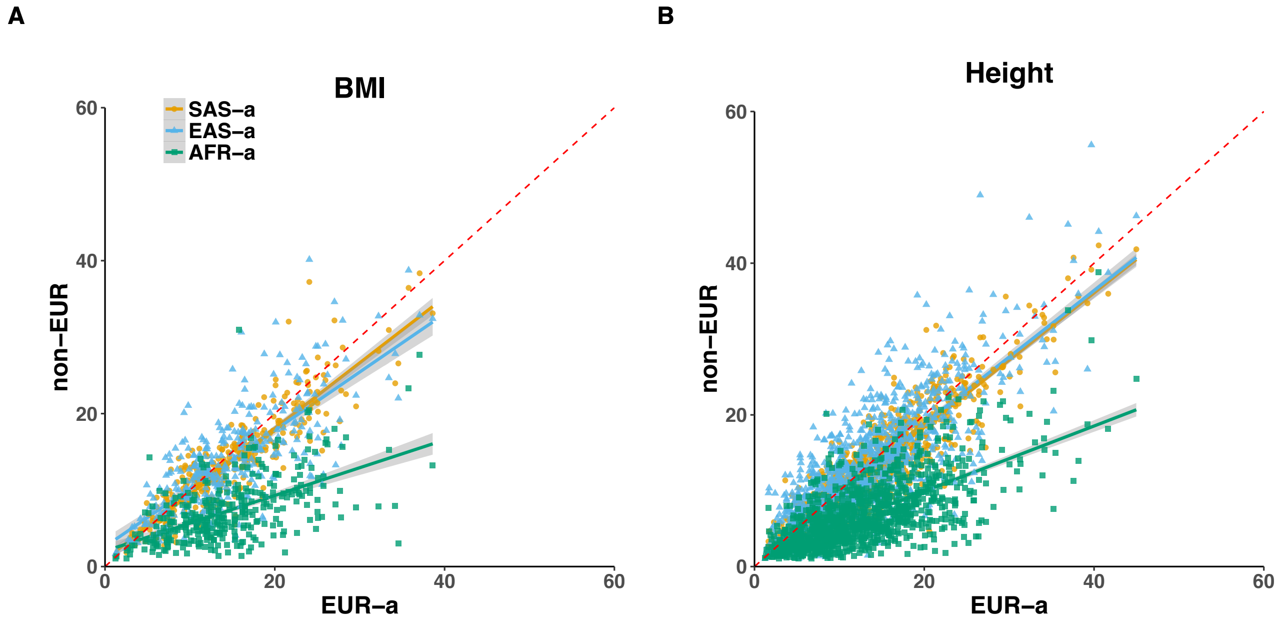


**Figure S7. The LD score correlation of trait-associated loci between ancestries.** We calculated the LD scores of those GWS SNPs for height and BMI, respectively (see details in **Supplementary Note 3**). 1KGP WGS data was used as the reference panel. We used a 100Kb window to calculate the LD scores and then compared them between the EUR-a and other non-EUR target populations (non-EUR). The red dashed line in each plot was: y=x.

**Table S1. Estimates of cross-population genetic correlations for height and BMI using HapMap3 SNPs.**

| Traits | Pops | $h^2_{\mathrm{EUR,SNP}}$(se) | $h^2_{\overline{\mathrm{EUR}},\mathrm{SNP}}$(se) | $\hat{r}_{g,\mathrm{SNP}}$(se) |
|---|---|---|---|---|
| | **EUR-a VS SAS-a** | 0.504(0.008) | 0.349(0.033) | 0.914(0.048) |
| **Height** | **EUR-a VS EAS-a** | 0.504(0.008) | 0.397(0.123) | 0.700(0.120) |
| | **EUR-a VS AFR-a** | 0.503(0.008) | 0.319(0.053) | 0.783(0.079) |
| | **EUR-a VS SAS-a** | 0.243(0.008) | 0.227(0.035) | 0.829(0.077) |
| **BMI** | **EUR-a VS EAS-a** | 0.242(0.008) | 0.185(0.122) | 0.776(0.278) |
| | **EUR-a VS AFR-a** | 0.242(0.008) | 0.181(0.055) | 0.774(0.143) |

$h^2_{\mathrm{EUR,SNP}}$ and $h^2_{\overline{\mathrm{EUR}},\mathrm{SNP}}$ refer to the SNP heritability calculated based on HapMap3 in Population 1 and Population 2, respectively. $\hat{r}_{g,\mathrm{SNP}}$ is the joint-SNP effects correlation between ancestries. The details are shown in **Supplementary Note 7**.
